# Comprehensive deletion scan of anti-CRISPR AcrIIA4 reveals essential and dispensable domains for Cas9 inhibition

**DOI:** 10.1101/2024.07.09.602757

**Authors:** Annette B Iturralde, Cory A Weller, Simone M Giovanetti, Meru J Sadhu

## Abstract

Delineating a protein’s essential and dispensable domains provides critical insight into how it carries out its function. Here, we developed a high-throughput method to synthesize and test the functionality of all possible in-frame and continuous deletions in a gene of interest, enabling rapid and unbiased determination of protein domain importance. Our approach generates precise deletions using a CRISPR library framework that is free from constraints of gRNA target site availability and efficacy. We applied our method to AcrIIA4, a phage-encoded anti-CRISPR protein that robustly inhibits SpCas9. Extensive structural characterization has shown that AcrIIA4 physically occupies the DNA-binding interfaces of several SpCas9 domains; nonetheless, the importance of each AcrIIA4 interaction for SpCas9 inhibition is unknown. We used our approach to determine the essential and dispensable regions of AcrIIA4. Surprisingly, not all contacts with SpCas9 were required, and in particular, we found that the AcrIIA4 loop that inserts into SpCas9’s RuvC catalytic domain can be deleted. Our results show that AcrIIA4 inhibits SpCas9 primarily by blocking PAM binding, and that its interaction with the SpCas9 catalytic domain is inessential.

**Significance:** For decades, researchers have determined the functionally important parts of proteins by deleting protein segments and seeing which ones affect protein function. This provides critical information about how the protein works. Here, we developed a high-throughput method to generate and test deletions in a protein of interest. Our method uses a CRISPR library approach, and can generate thousands of precisely programmed deletions in a single protein. We used it to test the effects of deletions in the phage anti-CRISPR protein AcrIIA4. Previous studies have shown that AcrIIA4 binds Cas9 using several interfaces, but the individual importance of these interfaces was unclear. We find that AcrIIA4 acts by blocking PAM binding, while its interaction with Cas9’s catalytic domain is dispensable.

## Introduction

CRISPR systems consist of RNA-guided endonucleases that cleave nucleic acid sequences specified by the variable spacer sequence of a bound guide RNA (gRNA) (1). CRISPR systems are widespread in bacteria and archaea, where they can provide adaptive immunity against viruses, plasmids, and other mobile genetic elements (MGEs) through gRNAs that recognize foreign DNA or RNA and target their cleavage. The adaptability of CRISPR endonucleases to cleave specific DNA sequences or otherwise recruit proteins to specific genomic sites is used in myriad ways in molecular biology research, predominantly featuring the type IIA Cas9 protein from *Streptococcus pyogenes*, referred to as SpCas9. gRNA-bound SpCas9 cleaves double-stranded DNA if its sequence matches the gRNA’s 20-nucleotide (nt) spacer sequence followed by the protospacer motif (PAM) sequence of NGG.

To counter the native role of CRISPR in host defense, MGEs encode a variety of anti-CRISPR proteins, or Acrs (2). Acrs have diverse strategies to block CRISPR cleavage, including through disruption of gRNA function, DNA binding, or DNA cleavage. Acrs have been repurposed technologically to regulate CRISPR, including to restrict CRISPR activity to a particular tissue type or to reduce cytotoxicity associated with CRISPR (3, 4). One of the most well-characterized Acrs is AcrIIA4, a potent inhibitor of SpCas9 identified from a *Listeria monocytogenes* prophage (5). AcrIIA4 is a compact 87-amino acid protein composed of a central three-stranded β-sheet flanked by three α-helices (6–8), as well as a 3_10_ helix in the β1-β2 loop that deforms upon SpCas9 binding (9). AcrIIA4 occupies the DNA-binding interfaces of several domains of SpCas9 (Fig. 1A). AcrIIA4 binds the SpCas9 C-terminal domain (CTD), which binds the PAM sequence of target DNA, using its β3 strand, α3 helix, and β2-β3, β3-α2, and α2-α3 loops. Residues in AcrIIA4’s α1-β1 and β2-β3 loops interact with the SpCas9 topoisomerase-homology (TOPO) domain, which aids in PAM binding. Finally, the β1-β2 loop of AcrIIA4 occupies the active site of SpCas9’s RuvC domain, which cleaves the nontarget DNA strand. However, the individual importance of these interactions to SpCas9 inhibition has not been determined.

**Figure 1.**
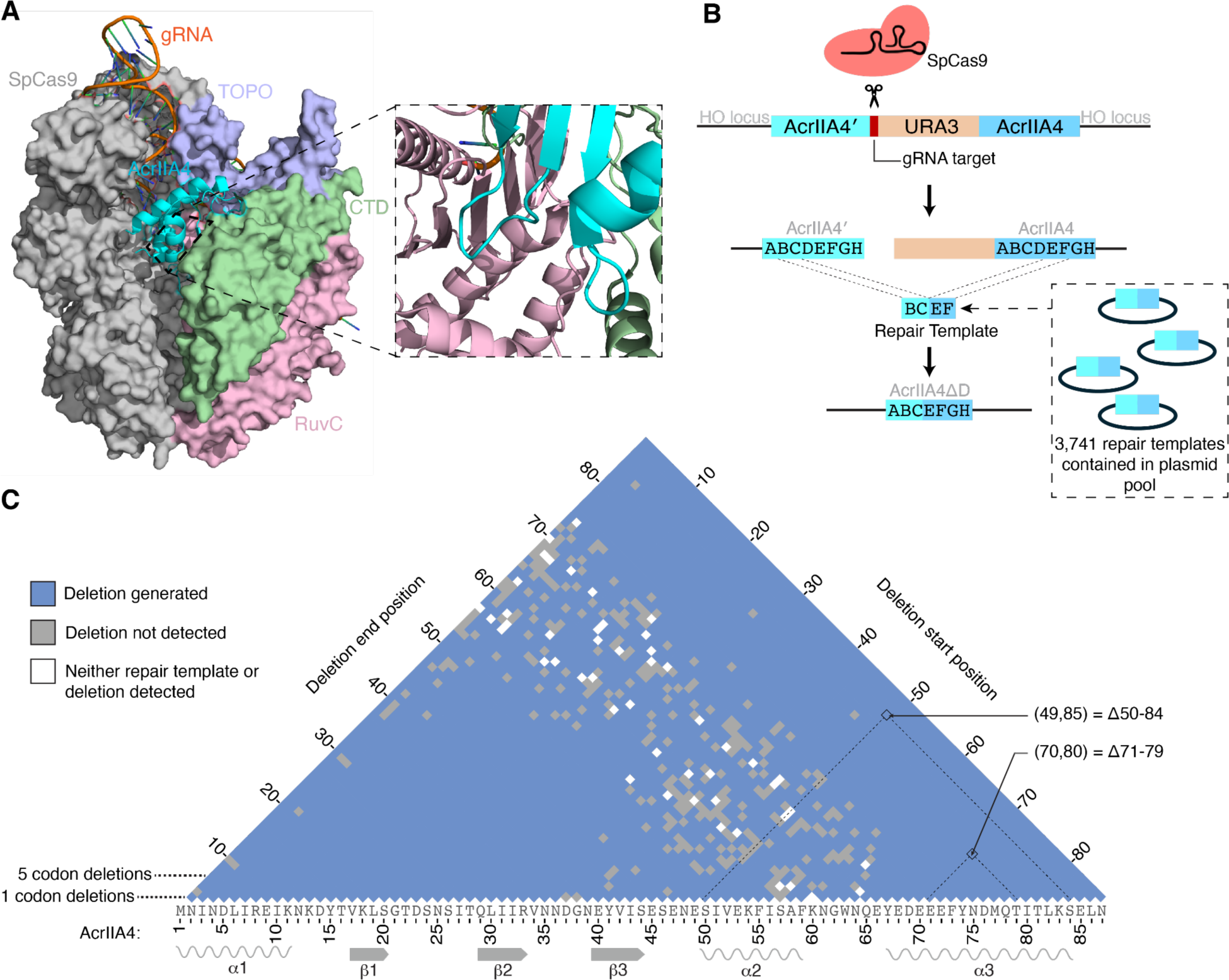
Comprehensive generation of in-frame deletions in AcrIIA4, an inhibitor of SpCas9. (A) Structure of AcrIIA4 bound to SpCas9 (PDB 5VW1) (7). AcrIIA4 (cyan) binds SpCas9’s CTD (green), TOPO (lavender), and RuvC (pink) domains. The inset shows the AcrIIA4-RuvC binding interface, which is largely obscured in the larger view by CTD. (B) Strategy for generating all possible in-frame deletions in AcrIIA4 in yeast. CRISPR cleavage using a constant gRNA is followed by repair directed by one of 3,741 deletion-generating repair templates. (C) High-throughput sequencing of AcrIIA4 alleles following deletion introduction identified 87% of the possible in-frame deletions, spanning the range of possible deletion sizes.

To precisely determine which domains of AcrIIA4 are essential for anti-CRISPR function, we generated nearly all possible in-frame deletions of all sizes in AcrIIA4 and tested their impact on SpCas9 inhibition. Functional domains of proteins have been mapped for decades by determining the effects of deleting segments of the protein (10). Augmenting such approaches to assay all possible in-frame deletions enables full and precise delineation of a protein’s essential and dispensable domains in an unbiased fashion (11), but creating all possible sizes of deletions across a gene scales quadratically with gene size and quickly amounts to thousands of deletions, necessitating a high-throughput approach. CRISPR technology can be used for high-throughput genetic interrogation, by synthesizing a library of DNA molecules encoding gRNAs and/or repair templates and using it to generate genetic variants in a pool of cells that contains CRISPR machinery (12–14). Here, we developed a high-throughput pooled CRISPR approach to comprehensively determine the effects of deletions in a protein and applied it to AcrIIA4. We find that the AcrIIA4 regions responsible for blocking the PAM-interacting domains of SpCas9 are essential, whereas the segment involved in blocking the RuvC catalytic domain is dispensable.

## Results

We developed an approach for comprehensive deletion mutagenesis that uses CRISPR in the budding yeast *Saccharomyces cerevisiae* (Fig. 1B). We used synonymous variants to generate two alleles of *acrIIA4* with maximum DNA sequence divergence while reflecting yeast codon usage, resulting in 61% DNA identity. We inserted the two *acrIIA4* alleles in tandem into the yeast genome at the *HO* locus, placing them after a galactose-inducible promoter and placing a synthetic Cas9 target site and the *URA3* gene from *Kluyveromyces lactis* between them. To generate an *acrIIA4* allele carrying a deletion between any pair of codons *n*_1_ and *n*_2_, we direct CRISPR cleavage at the synthetic target site between the *acrIIA4* copies and provide a repair template with homology to the first acrIIA4 copy up to codon *n*_1_, followed by homology to the second copy starting from codon *n*_2_. Thus, the use of this repair template during homology-directed repair produces a single fusion *acrIIA4* gene that has the region from codon *n*_1_ to codon *n*_2_ deleted. The fusion event also deletes the CRISPR target site and *URA3*, precluding further cutting and allowing for the selection of cells with successful deletions through their ability to grow on media containing 5-fluoroorotic acid (5-FOA), which is toxic to *S. cerevisiae* cells expressing *URA3* (15).

There are 3,741 possible unique in-frame *acrIIA4* deletions. To generate a comprehensive library of deletions we used large-scale array-based synthesis to synthesize a pool of all possible deletion-directing repair templates, composed of 100 nt matching the first *acrIIA4* copy followed by 100 nt matching the second. We cloned the repair template oligos into three replicate 2-micron plasmid libraries for high-copy maintenance in yeast cells. Massively parallel sequencing of the repair templates from two of the resulting plasmid libraries revealed 3,573 (95%) repair templates were represented in at least one of the replicate plasmid pools. The plasmid pools also encoded a gRNA targeting the novel sequence inserted between the *acrIIA4* genes, which was designed to have at least six mismatches with any other SpCas9 target site in the *S. cerevisiae* genome. Note that as the gRNA target site is constant, this approach is not constrained by the density of gRNA target sites in the *acrIIA4* gene, nor is there any aspect of variability in gRNA efficacy.

We transformed the repair template library into the *acrIIA4-URA3-acrIIA4* yeast strain. The strain also carried two copies of SpCas9, one under a *MAL62* promoter and the other under a *GAL*L promoter, allowing SpCas9 to be activated by either maltose or galactose (16–18). Both promoters are repressed by glucose. Transformed cells were selected on glucose plates and then re-plated on maltose to induce the maltose-responsive SpCas9 to generate deletion-carrying *acrIIA4* alleles. Resulting colonies were then grown on 5-FOA plates to select cells that had deleted *URA3* and thus were carrying the designed *acrIIA4* deletion alleles.

We sequenced the *acrIIA4* alleles present in the 5-FOA-resistant population by massively parallel sequencing. We recovered 3,305 distinct programmed deletions across the three replicates, 88% of the total possible deletions (Fig. 1C). Deletions were recovered across the entire range of sizes.

Next, we determined the ability of the *acrIIA4* deletion alleles to inhibit SpCas9 activity. We generated a plasmid expressing a gRNA that robustly targets the essential gene *ARB1* (14); usage of this gRNA for CRISPR cutting is lethal. To identify *acrIIA4* deletion alleles that maintained anti-CRISPR function, we transformed the *ARB1*-targeting gRNA plasmid into our pool of cells with programmed *acrIIA4* deletions and induced SpCas9 and *acrIIA4* expression on galactose plates. Approximately 30% as many colonies survived CRISPR induction compared to glucose plates without CRISPR expression. We used massively parallel sequencing of the *acrIIA4* genomic locus to identify the *acrIIA4* alleles present in survivors. Read counts were normalized against the count of each allele from the uninduced glucose condition (Fig. 2A).

**Figure 2.**
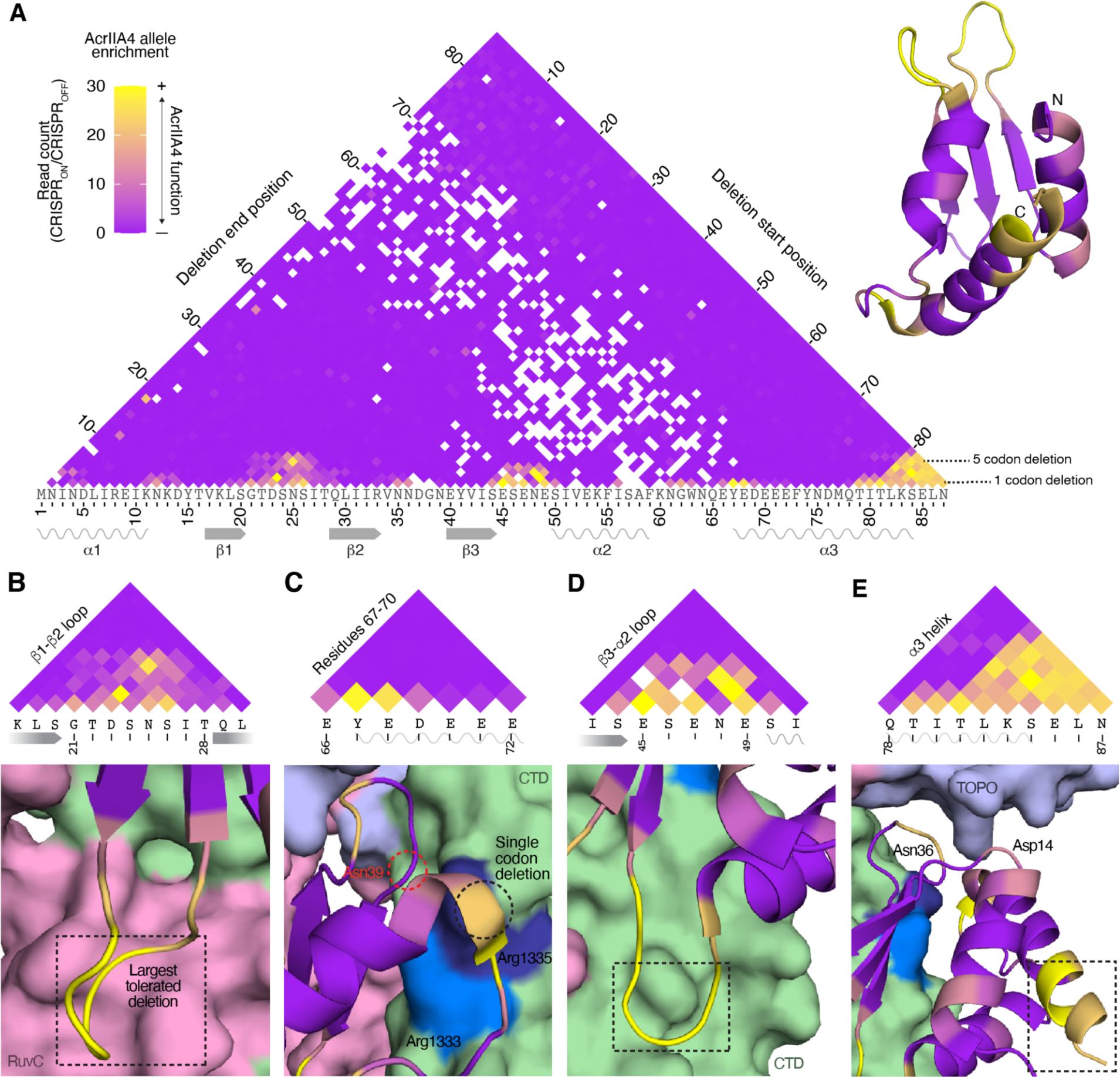
Functional characterization of AcrIIA4 deletion alleles. (A) Functionality of AcrIIA4 deletion alleles, determined by their read counts in CRISPR_ON_ conditions normalized against CRISPR_OFF_ read counts. Deletions are colored by their median value across three replicate experiments to minimize the effects of outlier escaper mutants; alleles only present in two replicates are represented by the lower of the two values. Residues in the AcrIIA4 crystal structure (right) are colored by the maximum functionality of deletions encompassing that residue (six-residue deletions or shorter). (B-E) Dashed boxes demarcate the largest tolerated AcrIIA4 deletion in the view. (B) Large dispensable regions within the RuvC-interacting β1-β2 loop. (C) AcrIIA4 residues 67-70, which interact with Arg1333 and Arg1335, are tolerant of single amino acid deletions. Asn39, which interacts with Arg1335, is not dispensable. (D) Large dispensable regions within the CTD-interacting β3-α2 loop. (E) Asp14 and Asn36, TOPO-interacting residues, are partially dispensable. The end of the C-terminal helix α3 is dispensable; note that the final two amino acids are not resolved in the crystal structure but were also dispensable.

We observed three distinct clusters of AcrIIA4 deletions that maintain function, in the β1-β2 loop, the β3-α2 loop, and the end of the α3 helix (Fig. 2A, Supplementary Fig. 1). Given the contacts between the β1-β2 loop and the catalytic site of the RuvC domain, the loop’s tolerance of deletions was notable. The loop is eight amino acids long; we found that AcrIIA4 retained anti-CRISPR function even when six of these amino acids were deleted (Fig. 2B). As this comprises the major RuvC-interacting surface, we propose that RuvC binding is dispensable for AcrIIA4’s anti-CRISPR function.

AcrIIA4 makes contacts with the SpCas9 CTD through residues in the β3 strand, α3 helix, and β2-β3, β3-α2, and α2-α3 loops. In CRISPR targeting, arginines Arg1333 and Arg1335 of the SpCas9 CTD engage the two guanines in the NGG PAM sequence.

Residues 67-70 in the α2-α3 loop and α3 helix of AcrIIA4, which includes three negatively charged residues, interact with the positively charged arginines (6). Interestingly, this segment of AcrIIA4 tolerated single amino-acid deletions, but not any larger deletions (Fig. 2C). We propose that this interaction is required for SpCas9 inhibition and can tolerate the removal of a single negative charge from AcrIIA4, but no more. Supporting the importance of this interaction, we see that Asn39 in the β2-β3 loop, which also contacts Arg1335, was not dispensable. In contrast, the β3-α2 loop, which contacts a distal component of the SpCas9 CTD, was a hotspot for tolerated deletions and could be shortened by up to four amino acids (Fig. 2D). This suggests that the critical CTD contacts are precisely the ones at the PAM-interacting interface.

AcrIIA4 makes relatively fewer contacts to SpCas9’s TOPO domain, which is also involved in PAM binding. TOPO residues that interact with the PAM are contacted by the AcrIIA4 residues Asp14 and Asn36 (6), both of which were partially dispensable (Fig. 2E). We noted that Asn36 is adjacent to Asn35 in the β2-β3 loop, and while each asparagine was individually partially dispensable, simultaneous deletion of both was not tolerated. Thus, binding of TOPO by AcrIIA4 appears to be important for AcrIIA4 function, with functional redundancy buffering the loss of individual contacts.

We observed a large number of tolerated deletions towards the end of the C-terminal helix α3 (Fig. 2E), with up to six terminal residues being dispensable. This segment does not make any contacts with SpCas9 and is largely separate from the rest of AcrIIA4. Consistent with the C-terminus being mutable, a C-terminal GFP fusion does not disrupt anti-CRISPR activity (19). In contrast, deletions of seven or more terminal residues were not tolerated. These deletions remove Ile80, which contributes to the hydrophobic core of AcrIIA4 along with Leu6, Ile10, and Phe55. Intriguingly, we observed that three six-residue deletions encompassing Ile80 retained partial function in all three replicates: Δ78-83, Δ79-84, and Δ80-85 (Supplementary Fig. 2). These deletions move Leu86 to position 80, which we propose partially reconstitutes Ile80’s role in AcrIIA4 stability.

We were curious whether the dispensable segments we identified in AcrIIA4 were ever absent in naturally occurring AcrIIA4 homologs. We identified 124 homologs of AcrIIA4 through a BLASTp search of the NCBI sequence database (20). Aligning these to AcrIIA4 revealed that 48 had gaps relative to AcrIIA4, falling into six unique patterns (Fig. 3A). Strikingly, most observed alignment gaps were found in regions we identified as dispensable. These included gaps in the β1-β2 and β3-α2 loops and truncations of the C-terminus. The largest gap was seen in several homologs that had a β1-β2 loop five amino acids shorter than in AcrIIA4, consistent with our observation that the AcrIIA4 β1-β2 loop could be shortened by up to six amino acids while maintaining anti-CRISPR function.

**Figure 3.**
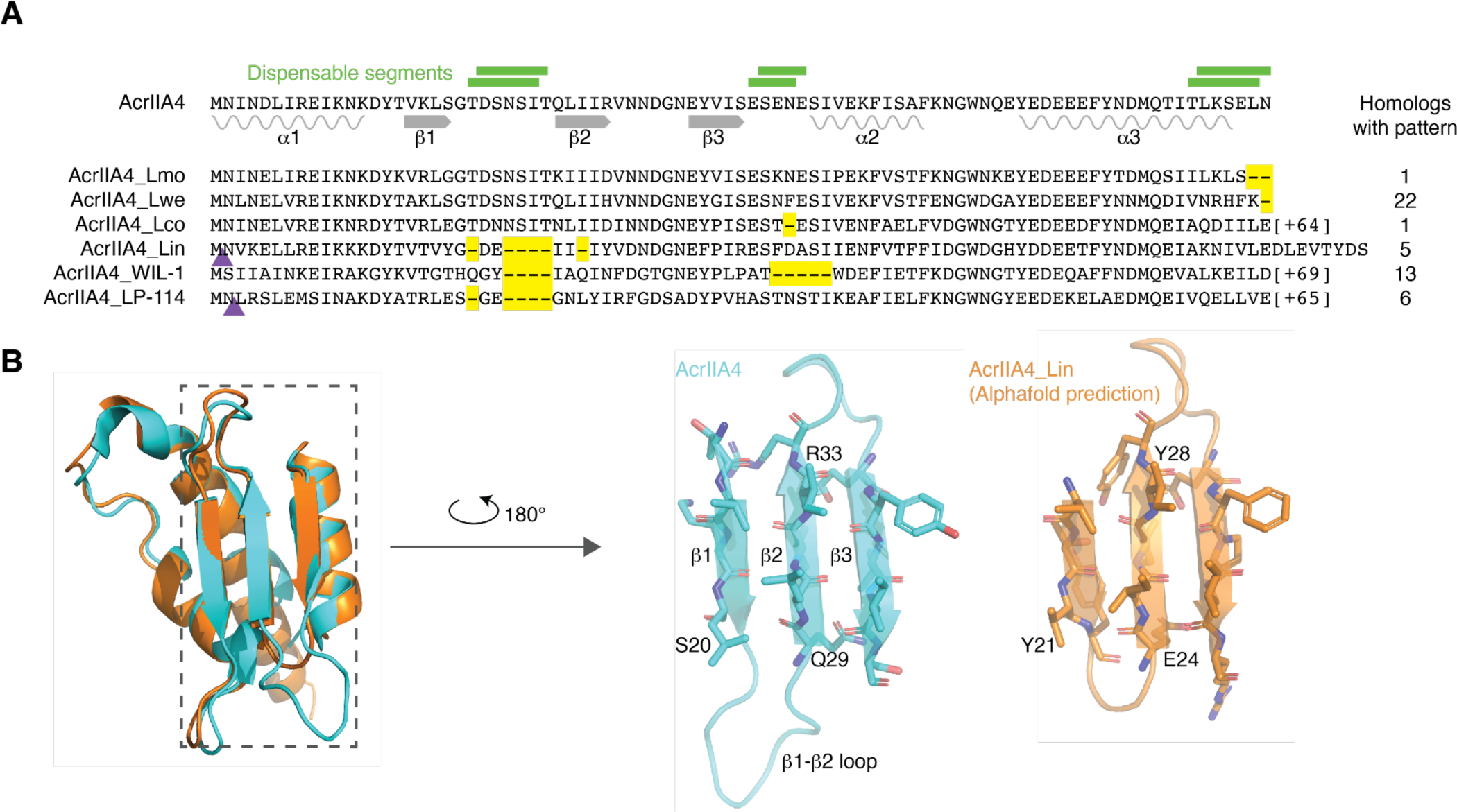
Deletions in naturally observed homologs of AcrIIA4 occur in experimentally determined dispensable segments. (A) Alignment of six AcrIIA4 homologs representing the identified deletion patterns. Yellow dashes signify deletions relative to AcrIIA4, while purple triangles demarcate the positions of insertions. The positions of select segments identified as dispensable in Fig. 2 are illustrated with green lines above the AcrIIA4 sequence. From top to bottom, the AcrIIA4 homolog NCBI IDs and source are EAD0751510.1 isolated from *L. monocytogenes*; MBC1462049.1 from *L. welshimeri*, WP_185587529.1 from *L. cossartiae*, WP_185342258.1 from *L. innocua*, YP_010843666.1 from phage WIL-1, and YP_009045111.1 from phage LP-114. (B) The AlphaFold-predicted structure of AcrIIA4_Lin has a full-length β2 strand.

One alignment gap was present in a non-dispensable segment in the AcrIIA4 homolog from *Listeria innocua* (NCBI ID WP_185342258.1, hereafter referred to as AcrIIA4_Lin_), which appeared to have a one-amino acid shortening of the indispensable C-terminal end of the β2 strand, in addition to a five-amino acid shortening of the dispensable β1-β2 loop. We generated an AlphaFold model (21) of the AcrIIA4_Lin_ protein structure (Fig. 3B) and found that the β2 strand in the model was five amino acids long, the same length as in AcrIIA4, while the modeled β1-β2 loop in AcrIIA4_Lin_ was a full six amino acids shorter than in AcrIIA4, consistent with our observed deletion tolerances.

We compared our results to three computational approaches for predicting the effects of deletions in AcrIIA4. First, we used AlphaFold to predict structures for all AcrIIA4 deletions of 10 amino acids or fewer, and then used FoldX to calculate the differences in protein stability between the predicted deletion-containing structures and the wild-type structure (22). We found many similarities between the predicted stability effects and our results, such as deletions being more tolerated in loops and at the C-terminus, and the tolerance of the 6-amino acid deletions that replaced Ile80 with Leu86 (Fig. 4A, B). The main departure was that many more large deletions were predicted to be stable than we found to be functional, especially in the β1 strand and the β2-β3 and α2-α3 loops. This could suggest these deletions form stable but nonfunctional proteins, or it could reflect bias in AlphaFold towards predicting structures to be stable (23). Next, we predicted AcrIIA4 deletion functionality using ESM1b, which predicts a sequence’s likelihood to exist given the “language” of known protein sequences (24). The functionality of a variant can be predicted by comparing the likelihood of the variant sequence to the likelihood of the wild-type sequence. For deletions of single amino acids in AcrIIA4, ESM1b also generally predicted higher deletion tolerance in loops and at the C-terminus (Fig. 4C), though it did not match the experimental data as closely as the AlphaFold-based stability predictions did. The performance of ESM1b decreased for larger deletions, which it often predicted would be more likely than the wild-type AcrIIA4 sequence. Also, no deletions had scores lower than −7, which is the recommended cutoff for calling a variant deleterious (24). Perhaps the poor performance of ESM1b stems from AcrIIA4 being a small protein that does not have homology to any described protein domains, which may lead to its sequence not following patterns learned from other protein sequences. Last, we used PROVEAN, which predicts the impact of protein variants by their consequences on alignments to homologous protein sequences (25).

**Figure 4.**
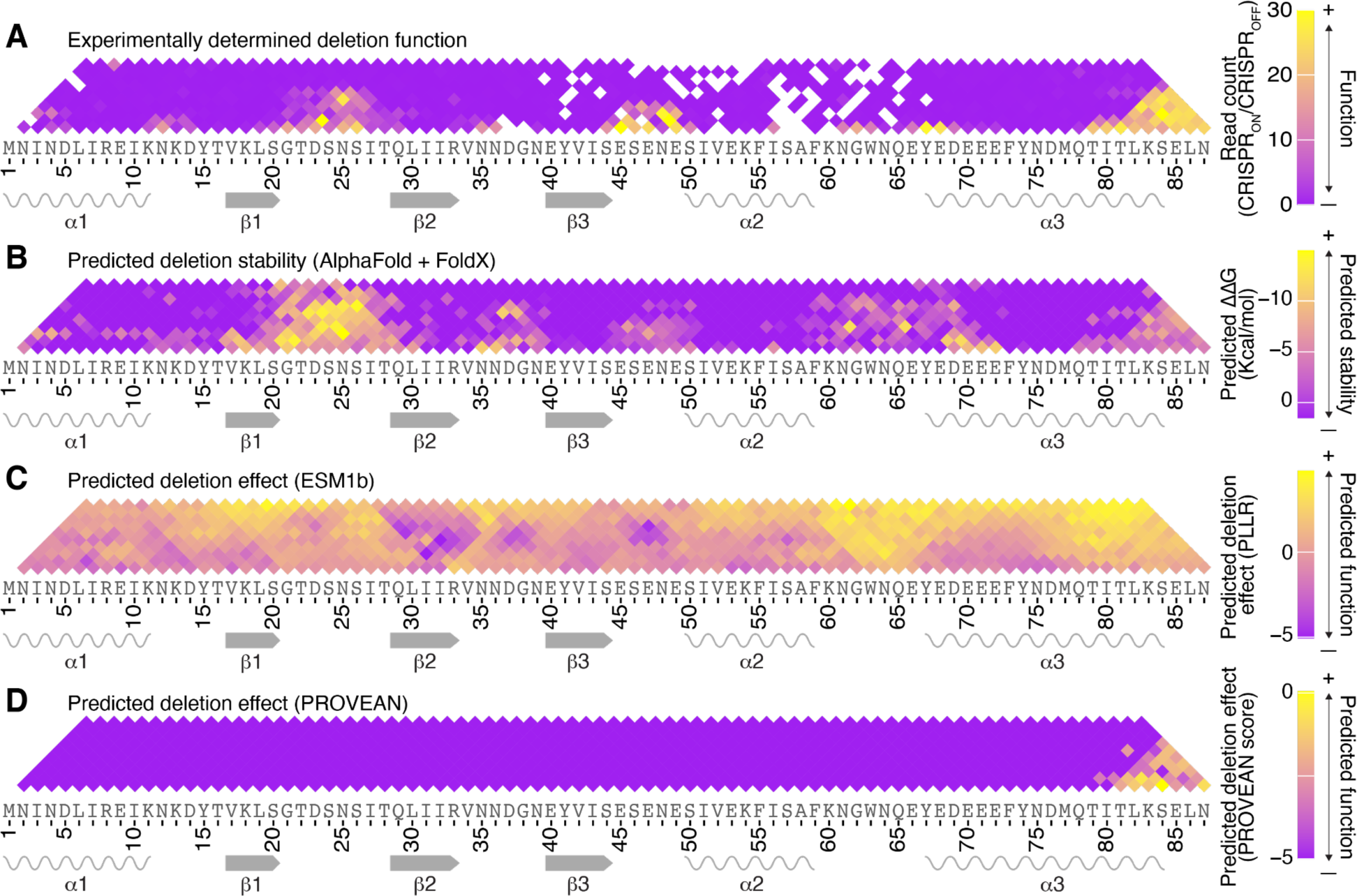
Predictions of the effect of deletions of 10 amino acids or fewer in AcrIIA4. (A) Our experimentally determined deletion effects. (B) The effect of deletions on AcrIIA4 stability predicted using AlphaFold2 to predict protein structures and FoldX to calculate the Gibbs free energy of folding, and then determining the change in free energy from the wild-type AcrIIA4 prediction. (C) Deletion effects predicted by ESM1b, calculated as the pseudo-log likelihood ratio (PLLR) between the wild-type and deletion variants. (D) Deletion effects predicted by PROVEAN.

Using a PROVEAN score threshold of −5 for loss-of-function, all deletions were predicted to be deleterious aside from those at the C-terminus (Fig. 4D). A more permissive threshold of −10 predicted functionality for some additional small deletions, often in loops but also in the α1 helix and the β strands (Supplementary Fig. 3C). We note that the recommended PROVEAN score threshold for loss-of-function is −2.5 (26). Overall, we found that the three computational predictors of deletion effect had different sensitivities and accuracies, and that predicting the effects of larger deletions is a particular challenge. None of the three predictors captured the increased dispensability of the RuvC-interacting residues relative to the residues that contact Cas9’s PAM-interacting domains.

## Discussion

AcrIIA4 is a robust inhibitor of SpCas9. It physically interacts with SpCas9 through an extensive interface that involves several distinct SpCas9 domains, including domains involved in PAM recognition (CTD and TOPO) and DNA cleavage (RuvC), though the importance of each individual interaction to SpCas9 inhibition was unknown. To elucidate the mechanism of AcrIIA4 inhibition of SpCas9, we comprehensively deleted segments of AcrIIA4 and assayed their function. We found that the interaction with the RuvC domain, mediated by the β1-β2 loop of AcrIIA4, is dispensable, as this eight-amino acid loop could be shortened by up to six amino acids without loss of anti-CRISPR activity. Our finding that AcrIIA4 does not need to occupy the RuvC active site to inhibit SpCas9 is consistent with the fact that AcrIIA4 robustly inhibits catalytically inactive Cas9 from binding DNA, which necessarily precedes DNA cleavage (8). Still, it remained possible that AcrIIA4 binding to the RuvC domain contributed to the binding energy required to block DNA binding. Based on our findings, binding to Cas9’s PAM-interacting domains is sufficient.

Though the physical interactions between AcrIIA4 and SpCas9 are largely mediated by the loops between secondary structural elements, we observed that none of the α helices and β strands of AcrIIA4 were fully dispensable. This likely reflects intramolecular interactions necessary for AcrIIA4 stability and functional shape, matching the observation that evolutionarily observed indels rarely occur in α helices and β strands (27). Consistent with this hypothesis, we observed that partial deletions of helices α1 and α3 were tolerated. These helices are at the very start and end of the protein, and the tolerated deletions were not near the AcrIIA4 core, so deletion of these segments might be less likely to disrupt AcrIIA4’s structure. Indeed, AlphaFold-predicted structures of these deletions were calculated to have comparable stability to the WT structure (Fig. 4B). Deletion scans of other proteins have similarly found higher tolerance for deletions at the N- and C-termini of domains (28).

Interestingly, we found several segments that could not be deleted despite being composed of smaller dispensable segments, as has been observed previously in other proteins (29). This could occur through functional redundancy, as we propose for amino acids 67-70, which has several negatively charged residues and can tolerate single amino acid deletions, but no further. Alternatively, a segment could have no direct role in protein function, but could link functional components that need to be a minimum distance apart. For instance, while multiple deletions of six amino acids were tolerated in the eight-amino acid β1-β2 loop, deletions of seven amino acids were not. We hypothesize that such deletions disrupt the relative positioning of strands β1 and β2, which may affect AcrIIA4 functionality while maintaining stability, based on AlphaFold’s stability predictions (Fig. 4B). The ubiquity of non-additive deletion effects highlights the utility of comprehensively testing deletions, which can reveal functional redundancy and dispensable segments that could be missed with a low-throughput approach.

A deep mutational scanning analysis of AcrIIA4 has been reported (30), measuring the ability of all possible nonsynonymous substitutions in AcrIIA4 to inhibit transcriptional repression mediated by catalytically inactive SpCas9 (CRISPRi). Though the two assays could have different sensitivities to Acr function, we see broad agreement in that positions that tolerated deletion also tolerated nonsynonymous variants, and positions that were intolerant of nonsynonymous variants did not tolerate deletion. However, positions that did not tolerate deletions often tolerated nonsynonymous variants, especially in α helices and β strands. This could reflect roles that secondary structures play in the structural integrity of AcrIIA4 or in the proper placement of the Cas9-interacting elements, as suggested above; such roles may be relatively indifferent to the biochemical properties of the secondary structures’ amino acid side chains. Another difference between these two analytical approaches is that deletion scanning additionally looks at the effects of deleting multiple amino acids at once, which allows for the analysis of functional redundancy and domain dispensability.

Comprehensive scanning of deletion effects has many applications in addition to mapping proteins’ essential and dispensable segments. It can also be used in genetic engineering to produce improved or novel protein function, as indels have been shown to have larger effects on protein function than missense variants (31). Similarly, deletions are an important source of evolutionary novelty (32, 33), so deletion scanning is useful to understand evolutionary trajectories. Deletion scanning can also be used in clinical genetics to characterize potential disease-causing deletion alleles.

Understanding the effects of deletion variants is essential for interpreting the risk of genetic disease from genome sequences: after single-nucleotide variants, deletions are the most common type of variant on ClinVar (34), and each individual person is predicted to harbor approximately 200 in-frame insertions or deletions relative to the human reference genome (35).

Our strategy for deletion scanning involved chimerizing two copies of *acrIIA4* following CRISPR cutting between them. A previous study used CRISPR to generate cuts within a single copy of yeast *SGS1*, which were repaired using repair templates that directed 20-amino acid deletions across the gene body (36). By cutting between two copies of our target gene, we did not require PAM sequences close to the regions being deleted, allowing us to comprehensively characterize deletions of small to large size. In addition, our strategy uses a constant gRNA, which avoids issues arising from variable gRNA targeting efficiency. Finally, by concurrently deleting a *URA3* gene between the target gene copies, we could select for cells carrying deletion alleles.

Deletion scans have also been carried out using non-CRISPR approaches. One strategy is to use oligonucleotide synthesis technology to directly generate deletion alleles, rather than to generate repair templates that direct deletions following CRISPR cutting. As oligonucleotides synthesized economically at large scale are currently limited to lengths of 300 nt or fewer, to date this approach has been applied to short genes, segments of genes, or small deletions within larger genes (11, 28, 29, 33). Another strategy is to introduce restriction sites into the gene of interest, cleave, and then self-anneal to produce truncated gene products. For deletion scanning methods that use this framework (31, 37), a challenge is that novel sequences are inserted at the deletion junction for most or all of the deletion alleles.

An inherent challenge to the concept of deletion scanning is that the number of potential deletions scales quadratically with the size of the protein, which could theoretically be impractical for a large gene even when using a high-throughput approach. We note that the oligonucleotide library synthesis product we used can produce up to 244,000 sequences in a single order. With our approach, that would allow comprehensive deletion scans of proteins up to 700 residues, encompassing most proteins. More importantly, our approach allows specific deletions to be programmed, such as deletions under a certain size or deletions within predicted domains or secondary structure elements. Screening select deletions would lead to a linear scaling with protein length and thus prevent the number of tested deletions from becoming unmanageable.

We observed that the repair template plasmid pool had lower representation of repair templates targeting deletions that ended around codons 50-70 of acrIIA4 (Supplementary Fig. 4A), leading to a moderate depletion of those deletions from our set of recovered deletion alleles (Fig. 1C). We suspect that the DNA sequence around codons 50-70 of the second acrIIA4 copy is challenging either for the oligonucleotide synthesis or for the library cloning. The representation of deletions in this range was sufficient for our analyses, but if a more complete representation of deletions in that range had been desired, we would expect it to be possible through targeted amplification of those repair templates from the oligonucleotide library, resynthesis of that subset of oligonucleotides, or re-coding of that region using synonymous codons. Indeed, we were able to fully remedy a similar issue that arose in our initial cloning using synonymous re-coding of a segment of the first acrIIA4 copy (see Methods).

For applications other than mapping essential domains, such as determining the evolutionary or pathogenic potential of genetic variants, it can be interesting to test insertions as well as deletions (11, 33). Our method could be easily adapted to additionally generate internal duplicated segments, which are the most common form of naturally occurring insertion mutations (38), by having repair templates in which the start point in the second gene copy comes before the end point of the first copy.

Homologous proteins often have divergent functions, and chimeras between them can map the causative sequence differences. For instance, the yeast GTPases Sec4 and Ypt1 have different functions in the secretory pathway that were mapped to differences in loops 2 and 7 (39, 40). Chimeras have also been used to study orthologs between species, such as the human and macaque homologs of the innate immune protein TRIM5α, whose differences in HIV restriction were mapped to a loop in the SPRY domain (41–44). Our method could be readily adapted to generate chimeras between two different genes, rather than to make deletions. Generating and characterizing chimeras in high throughput would enable rapid fine-resolution determination of specific amino acids causing functional differences between homologs.

## Supporting information

Plasmid maps

Supplementary Table 2 - Plasmids

Supplementary Table 3 - Oligonucleotides

Supplementary Table 1 - Strains

## Acknowledgements

We thank Bryan Thurtle-Schmidt, Elena Zehr, two anonymous reviewers, and all members of the Sadhu lab for helpful discussion. We thank Jean-Marc Daran, Lars Steinmetz, and Leonid Kruglyak for strains and plasmids. This work utilized the computational resources provided by the National Institutes of Health (NIH) HPC Biowulf Cluster (http://hpc.nih.gov). This work was supported by the Intramural Research Program of the National Human Genome Research Institute, NIH (1ZIAHG200401).

## Author contributions

A.B.I., C.A.W., and M.J.S. conceptualized research; A.B.I., C.A.W., S.M.G., and M.J.S. performed formal analysis; A.B.I. performed investigation; A.B.I., C.A.W., and M.J.S. developed methodology; C.A.W. and M.J.S. performed supervision; A.B.I., C.A.W., S.M.G., and M.J.S. prepared visualizations; and A.B.I., C.A.W., and M.J.S. wrote the paper. All authors read and revised the paper.

## Methods

### Generation of base strains and plasmids

All strains, plasmids, primers, and oligonucleotides are included in Supplementary Tables 1-5.

All plasmids were transformed into NEB 5-alpha Competent *Escherichia coli* cells (New England Biolabs) and purified using Qiaprep Spin Miniprep Kits or Plasmid Plus Maxi Kits for plasmid libraries (QIAGEN). All PCR products were purified using QIAquick PCR Purification Kits or MinElute PCR Purification Kits (QIAGEN). Plasmid and genomic sequences were verified through Sanger sequencing (Eurofins Genomics, Louisville, KY, USA). All yeast transformations were done according to standard lithium acetate transformation procedures (45).

A plasmid carrying the two *acrIIA4* alleles was synthesized by Genewiz (MSp513). The sequence of the second *acrIIA4* allele matched the sequence used by Basgall, et al. (19). The sequence of the first *acrIIA4* allele was generated using custom python scripts in which each codon was selected to share as few bases as possible with the corresponding codon of the second *acrIIA4* allele. When multiple codons met that criterion, the codon was randomly selected from the available codons with weights matching codon usage in the *S. cerevisiae* genome. The first *acrIIA4* allele was preceded by the *GAL*L promoter (46) and the second was followed by the T_synth8_ terminator (47). The *acrIIA4* genes were flanked by 120-bp homology arms to the *HO* locus. This Acr construct was amplified using PfuUltraII fusion HS DNA Polymerase (Agilent Technologies) with the primers ANx87 and ANx88 and was transformed into strain MS262 (MSy15), which had *NEJ1* deleted to eliminate non-homologous end-joining as a repair pathways for DSBs (48) in favor of homology-directed repair (Supplementary Table 1). To increase transformation efficiency, during the transformation we directed a double-stranded DNA break at the *HO* locus by co-transforming the Acr construct with a plasmid constitutively expressing SpCas9 from the *TEF* promoter, acquired from Addgene (plasmid #: 162606; designated MSp181 in Supplementary Table 2), and a PCR construct encoding a gRNA targeting the yeast *HO* locus. The gRNA construct was generated by amplifying the targeting sequence from pMS58 (MSp20) with primers ANx84 and ANx85 (Supplementary Table 3). Positive transformants of the *acrIIA4* construct into the *HO* locus were selected by growth on plates lacking uracil and leucine (CSM-Ura-Leu; Sunrise Science Products) and confirmed by Sanger sequencing. A sequence-verified clone, in which MSp181 had been spontaneously lost, was designated MSy261.

MSy261 was used to perform the first deletion scan replicate experiment, as described in the section “Creation of acrIIA4 deletion alleles”. However, plasmids in the repair-template plasmid library (whose generation is described in the section “Design of repair templates”) that contained the sequence of codons 26-35 of the first copy of *acrIIA4* were strongly depleted from the plasmid library for unclear reasons, presumably related to that sequence posing a challenge either for oligonucleotide synthesis or library cloning. We thus generated a second strain with these codons re-coded, MSy262, which was used for the second and third deletion scan replicate experiments (Supplementary Fig. 4B,C). This was done by amplifying two segments of the *acrIIA4* construct on MSy261, upstream and downstream of codons 26-35, using Herculase II Fusion DNA Polymerase (Agilent) and introducing the alternate sequence by including it in the primers used for amplification (ANx80-83). To make MSy262, the two amplicons were transformed into MSy14, which carries a galactose-inducible SpCas9 plasmid, with the *HO*-targeting gRNA described above, and immediately plated onto CSM-Leu-Ura galactose plates.

Our strategy of introducing deletions into *acrIIA4* involved cutting with SpCas9 between the *acrIIA4* copies. For that step, we generated a maltose-inducible SpCas9 plasmid to induce SpCas9 expression while maintaining repression of the galactose-inducible *acrIIA4*, to prevent the anti-CRISPR activity of acrIIA4 from inhibiting the generation of deletions. We inserted a DNA construct containing the *MAL63* transcription factor gene and *MAL62* promoter sequence, ordered from Genewiz (MSp514), upstream of the SpCas9 gene on MSp14. We named the resulting plasmid MSp500, and it has been deposited on Addgene as plasmid # 227704. The sequence of the *MAL* genes was taken from the genome sequence of YJM145 (MSY28) (18). Amplification of the *MAL* locus sequence was done with KAPA Hi-Fi Hotstart DNA Polymerase (and primers ANx16.5 and 17.5), and inserted into MSp14 via Gibson assembly (New England Biolabs, Ipswich, MA, USA) following a digestion of the plasmid with SpeI-HF (New England Biolabs).

MSy261 was transformed with a plasmid carrying a galactose-inducible SpCas9, p415-GalL-Cas9-CYC1t (Addgene plasmid #:107922; MSp6) and transformants were selected on CSM-Ura-Leu glucose plates. One colony was picked and transformed with MSp500, with transformants selected on CSM-Ura-Leu + clonNAT (100ug/mL) glucose plates (MSy263). MSy262, already carrying a galactose-inducible SpCas9, was also transformed with MSp500 and plated on CSM-Ura-Leu + clonNAT (100ug/mL) glucose plates to produce MSy264.

### Design of repair templates

Custom python scripts were used to generate repair template sequences for directing all possible in-frame deletions within *acrIIA4*. In brief, to generate the repair template for the deletion from the *m*th to the *n*th codon, we appended the 100 nt up to the *m*th codon of the first *acrIIA4* allele to the 100 nt after the *n*th codon of the second *acrIIA4* allele.

For deletions close to the start of *acrIIA4*, their repair templates included sequence from the *GAL*L promoter preceding the first copy, and for deletions close to the end, their repair templates included the *acrIIA4* terminator and the sequence of the *HO* locus flanking the insertion site. We included the start codon and stop codon as deletable codons. We also designed repair templates that generated a clean chimera of the two *acrIIA4* alleles into a single *acrIIA4* without deletion, for 85 repair templates. Once all sequences were created, 15-nt primer sites were added to the beginning and end of the strings, making all sequences 230 characters long. Sequences were then ordered as an oligonucleotide pool through Agilent (Santa Clara, California, USA). This process was performed once to generate the repair templates used for replicate 1, and then again with the modified sequence of the first *acrIIA4* allele to generate the repair templates used for replicates 2 and 3 (Supplementary Tables 4 and 5). For the purpose of this experiment, we proceeded with data analysis of 3,741 sequences, excluding deletions of the start or stop codons, as well as repair templates that generated clean chimeras.

MSp218 was used as the backbone of the three replicate repair template plasmid pools, and was generated from pMEL16 (MSp50), a *HIS3* SpCas9 gRNA plasmid acquired from Jean-Marc Daran (Addgene plasmid #:107922). For cloning in the repair templates, we engineered EcoRI and BamHI sites downstream of the gRNA gene. In addition to carrying the repair templates, we designed MSp218 to encode a gRNA sequence targeting a sequence inserted between the two *acrIIA4* alleles, ACGTATCGGTACATGCGCAT. This sequence was picked to have equal representation of each of the four bases, as well as to have at least six mismatches to any other SpCas9 target sites in the yeast genome. MSp218 has been deposited on Addgene as plasmid # 227698.

We amplified the repair template oligonucleotide pools using primers that additionally added 12 nt of random DNA on each end to serve as barcodes as well as homology arms for Gibson assembly (ANx23 and 24); PCR was performed with PfuUltra II Fusion High-fidelity DNA Polymerase for 15 cycles. MSp218 was double digested using EcoRI-HF and BamHI-HF (New England Biolabs) on the cloning site, and the oligo amplicons were inserted via Gibson assembly. 50uL of Gibson assembly were transformed into 1000uL of 5-alpha competent *E. coli* cells for each replicate (New England Biolabs).

Replicate 1 produced 100,000 colonies, Replicate 2 produced 100,000 colonies, and Replicate 3 produced 80,000 colonies. Colonies were washed off plates and plasmid libraries were isolated with the Qiagen Plasmid Plus Maxi kit.

### Creation of *acrIIA4* deletion alleles

The repair template plasmid pools were transformed en masse (2.05 μg of Replicate 1; 3.5 μg of Replicates 2 and 3) into approximately 10^9^ cells of MSy263 or MSy264 as appropriate, and plated on CSM-Ura-Leu-His +clonNAT glucose plates. Replicate 1 produced 260,000 colonies, Replicate 2 produced 300,000 colonies, and Replicate 3 produced 250,000 colonies. Cells were then harvested by washing the plate with glucose-H,L,U +clonNAT liquid media, and a 100-mL culture of these cells was then started at an optical density (OD_600_) of 0.002 using the same media. The culture was left to grow overnight at 30°C until it reached an OD_600_ of 1. Approximately 2 × 10^9^ cells were then plated over multiple CSM Maltose His-Leu +cloNAT plates to induce SpCas9 from MSp500, resulting in 77,000 colonies for Replicate 1, 66,000 colonies for Replicate 2, and 80,000 colonies for Replicate 3. Cells were harvested by washing plates with YPD, and approximately 4 × 10^7^ cells were replated onto 5-fluoroorotic acid (5-FOA)-Leu-His plates. On 5-FOA plates, Replicate 1 produced 1.6 × 10^6^ colonies, Replicate 2 produced 1 × 10^6^ colonies, and Replicate 3 produced 1 × 10^6^ colonies.

### Characterization of *acrIIA4* deletion allele function

We generated a plasmid carrying a gRNA that directs CRISPR cleavage of the essential gene *ARB1* by inserting an *ARB1*-targeting gRNA sequence into pLK88 (MSp11), a *URA3* high-copy plasmid from Leonid Kruglyak. An oligonucleotide containing the gRNA sequence was ordered from Integrated DNA Technologies (ANx86) and inserted by Gibson assembly into MSp11 following a double digestion with SphI-HF and BstEII-HF (New England Biolabs). This plasmid was named MSp501 and has been deposited on Addgene as plasmid # 227705.

Yeast cells with *acrIIA4* deletion alleles were washed off 5-FOA plates using CSM-Leu-His glucose liquid media. A 50-mL culture was started at OD_600_ of 0.01 using the same liquid media, and it was left to grow overnight at 30°C. Once grown overnight, the culture was used to start a new culture in YPD starting at OD_600_ of 0.125, allowed to double twice, and then transformed with MSp501. In Replicate 1, 1.5 μg of MSp501 was transformed into approximately 10^9^ cells; in Replicates 2 and 3, 2 μg were transformed into approximately 10^9^ cells. Transformations were plated on CSM-Leu-His-Ura glucose plates, producing 300,000 colonies in Replicate 1, 600,000 colonies in Replicate 2, and 800,000 colonies in Replicate 3. Plates were then scraped once more and 1 × 10^6^ cells were plated on CSM-Leu-His-Ura galactose plates to simultaneously induce the *acrIIA4* deletion alleles and SpCas9 to target *ARB1*. An equal number of cells were plated on CSM-Leu-His-Ura glucose plates as controls. On glucose, Replicate 1 produced 1 × 10^6^ colonies, Replicate 2 produced 900,000 colonies, and Replicate 3 produced 1 × 10^6^colonies. On galactose, Replicate 1 produced 300,000 colonies, Replicate 2 produced 280,000 colonies, and Replicate 3 produced 370,000 colonies.

### Data collection and sequencing

Yeast genomic DNA was extracted from 4×10^7^ cells (per library) harvested off of the 5-FOA plates, galactose plates, and final glucose plates using the DNeasy Blood & Tissue Kit (QIAGEN). We then amplified the chimeric *acrIIA4* locus from the genomic DNA, using primers containing overhangs for attaching Illumina adapters. The forward primers for the genomic DNA annealed upstream of the first *acrIIA4* allele and the reverse primers annealed downstream of the second allele, such that they would amplify all chimeric *acrIIA4* alleles. Additionally, yeast plasmid DNA was extracted from 5×10^7^ cells harvested off of galactose and final glucose plates of replicates 2 and 3 using Qiaprep Spin Miniprep Kit (QIAGEN). We amplified the 200-bp repair template region from the initial *E. coli* plasmid pools of replicates 1 and 2 and the yeast plasmid extractions from replicates 2 and 3. Genomic primers (ANx25-34) and plasmid primers (ANx35-44) were staggered to maintain sequence diversity on the Illumina flow cell. All PCRs were performed using KAPA HiFi Hotstart ReadyMix DNA Polymerase for 20 cycles. A second PCR was performed to attach the Nextera adapter index primers using KAPA HiFi Hotstart ReadyMix DNA Polymerase for 10 cycles, after which samples were prepared for sequencing according to Illumina Miseq sequencing protocols.In total, four Miseq runs using Illumina Miseq 500 cycle V2 kits were performed: once for sequencing the genomic reads and *E. coli* plasmid pool of replicate 1, once for sequencing the *E. coli* plasmid pool of replicate 2, and once for sequencing the genomic and plasmid reads of replicates 2 and 3.

### Data analysis

PEAR software (Version 0.9.8) was used to process and align forward and reverse Miseq reads. Custom python and bash scripts were used to match Miseq sequencing reads of the genomic and plasmid DNA amplicons to the possible *acrIIA4* deletion alleles and generate read counts of perfectly matched reads to each deletion in each library. For the individual glucose and galactose genomic libraries, counts were first normalized by dividing the number of reads corresponding to a specific deletion by the total amount of reads in that library. We then divided all galactose values by their corresponding glucose value to calculate the enrichment of *acrIIA4* deletions in cells performing a lethal CRISPR edit. The analysis of deletion tolerance primarily utilized the read counts of deletion alleles amplified from the yeast genomic DNA. Analyzing replicate-replicate correlation plots between the three replicates, we found good agreement (r of 0.65, 0.68, and 0.68) when we capped gal/glu ratios at 30 (Supplementary Fig. 5A), which is slightly higher than the ratio for WT AcrIIA4. This indicated that variation above 30 was due to noise rather than biological significance, probably arising from variation in PCR amplification. We thus used this cap in downstream analyses. We also found higher correlations values between replicates for deletions of 10 or fewer residues (r of 0.79, 0.79, and 0.76) (Supplementary Fig. 5B), which we inferred was due to noise introduced by occasional spontaneous escaper mutants carrying large nonfunctional acrIIA4 deletions to high frequency. To minimize the effects of such escaper mutants, when merging the data from the three replicates for display in Fig. 2, we took the median of the galactose/glucose read count values for a given deletion across the replicates. A deletion was only counted as observed in a given replicate if it had at least five reads in glucose. If a deletion was not observed in one replicate, the lower of the remaining two values was taken. If there were no counts recorded for a given deletion in all three replicates, it was represented by a blank pixel on the heat map. For comparison, we also took the mean of the galactose/glucose read count values across replicates, and found that the dispensability of each analyzed domain was unaffected (Supplementary Fig. 6). We also performed a parallel analysis using the plasmid reads, and found that repair templates were enriched following CRISPR induction corresponding to the genomic AcrIIA4 deletion alleles we found to maintain anti-CRISPR function (Supplementary Fig. 7).

To determine the statistical significance of the domains that appeared to be dispensable (β1-β2 loop, β3-α2 loop, and the end of the α3 helix), we calculated the mean of the functional scores in the smallest triangular area encompassing the entirety of each domain, for each replicate. We then acquired the mean of an equal number of randomly selected data points as were contained in each triangular area and repeated this 1,000 times (Supplementary Fig. 1) for comparison by the Monte Carlo method.

To find naturally occurring AcrIIA4 homologs with deletions relative to the canonical AcrIIA4, we performed a BLASTp search on the NCBI BLASTp server using standard parameters. We used custom R scripts to identify homologs with deletions, and manually verified that the deletions did not result from incomplete sequencing.

ColabFold (version 1.5.5) (49), an open-source version of AlphaFold2 (50), was used to generate the structural models of AcrIIA4 homologs and deletion alleles through ChimeraX; of the predicted structures for each sequence, the highest confidence structure was used (restricting to relaxed structures for the predicted structures for deletion alleles). We then used the “stability” command in FoldX (Version 5.0) (22) to calculate the Gibbs free energy of folding (ΔG) for each deletion allele’s predicted structure. We subtracted the calculated ΔG for the wild-type AcrIIA4 structure from the ΔG value for each deletion to obtain ΔΔG values. We thresholded ΔΔG values at a maximum deleteriousness of 1.5, as that has previously been recommended as the threshold for destabilizing mutations (51). The analysis was not substantially changed when a much more permissive threshold of 15 was used (Supplementary Fig. 3B). For prediction of protein function using ESM1b, we used scripts from Brandes et al. for all deletions up to 10 amino acids in length (24). For prediction of protein function for all possible deletions using PROVEAN (25), we used the precise software version combinations of psiblast and blastdbcmd from ncbi-blast-2.4.0+, cd-hit version 4.8.1, PROVEAN version 1.1.5 and the RefSeq non-redundant database v4.

All statistical measurements and visualizations were done with custom R scripts (Versions 4.2.2 and 4.3.2) and R package ggplot2 (Versions 3.4.3 and 3.5.0) through RStudio. Crystal structure visualizations were generated with the PyMOL molecular visualization system, Versions 2.5.0 and Version 3.0.0 Open-Source, Schrödinger, LLC.

## Data Availability

- Python and R scripts: https://github.com/annetteig/AcrIIA4-Functional-Domains

